# High-throughput screening of the ReFRAME, Pandemic Box, and COVID Box drug repurposing libraries against SARS-CoV2 nsp15 endoribonuclease to identify small-molecule inhibitors of viral activity

**DOI:** 10.1101/2021.01.21.427657

**Authors:** Ryan Choi, Mowei Zhou, Roger Shek, Jesse W. Wilson, Logan Tillery, Justin K. Craig, Indraneel A. Salukhe, Sarah E. Hickson, Neeraj Kumar, Rhema M. James, Garry W Buchko, Ruilian Wu, Sydney Huff, Tu-Trinh Nguyen, Brett L. Hurst, Sara Cherry, Lynn K. Barrett, Jennifer L. Hyde, Wesley C. Van Voorhis

## Abstract

SARS-CoV-2 has caused a global pandemic, and has taken over 1.7 million lives as of mid-December, 2020. Although great progress has been made in the development of effective countermeasures, with several pharmaceutical companies approved or poised to deliver vaccines to market, there is still an unmet need of essential antiviral drugs with therapeutic impact for the treatment of moderate-to-severe COVID-19. Towards this goal, a high-throughput assay was used to screen SARS-CoV-2 nsp15 uracil-dependent endonuclease (endoU) function against 13 thousand compounds from drug and lead repurposing compound libraries. While over 80% of initial hit compounds were pan-assay inhibitory compounds, three hits were confirmed as nsp15 endoU inhibitors in the 1-20 μM range in vitro. Furthermore, Exebryl-1, a β-amyloid anti-aggregation molecule for Alzheimer’s therapy, was shown to have antiviral activity between 10 to 66 μM, in VERO, Caco-2, and Calu-3 cells. Although the inhibitory concentrations determined for Exebryl-1 exceed those recommended for therapeutic intervention, our findings show great promise for further optimization of Exebryl-1 as an nsp15 endoU inhibitor and as a SARS-CoV-2 antiviral.

**Author summary:** Drugs to treat COVID-19 are urgently needed. To address this, we searched libraries of drugs and drug-like molecules for inhibitors of an essential enzyme of the virus that causes COVID-19, SARS-CoV-2 nonstructural protein (nsp)15. We found several molecules that inhibited the nsp15 enzyme function and one was shown to be active in inhibiting the SARS-CoV-2 virus. This demonstrates that searching for SARS-CoV-2 nsp15 inhibitors can lead inhibitors of SARS-CoV-2, and thus therapeutics for COVID-19. We are currently working to see if these inhibitors could be turned into a drug to treat COVID-19.

## Introduction

Coronaviruses (CoVs) are a group of single-stranded positive sense RNA (+ssRNA) viruses belonging to the *Nidovirales* order. CoVs encode large RNA genomes (~30 kb) and infect diverse host species including humans, primate, bats, birds, and livestock. The ability of CoVs to jump host species has led to the emergence of multiple human coronaviruses, including the highly pathogenic SARS (severe acute respiratory syndrome) and MERS (middle eastern respiratory syndrome) viruses. The recent emergence of SARS-CoV-2 and the subsequent COVID-19 pandemic has already resulted in 1.7 million deaths worldwide as of mid-December, 2020, and is projected to cause over 2.8 million deaths by April 2021 despite the recent deployment of multiple vaccines (https://covid19.healthdata.org/global). Despite the recent approval of two SARS-CoV-2 vaccines, which promises to turn the tide of the current pandemic, it will be some time before a significant proportion of the population can be vaccinated in order to achieve the level of community protection necessary in the community and to prevent the spread of the virus. Furthermore, recent reports of emerging strains with different pathogenic properties have raised concerns that the current vaccines may prove less efficacious over time as new variants emerge. Therefore, there continues to be significant need for the development of small molecule inhibitors to complement existing therapeutics, not only for SARS-CoV-2, but also for related beta-coronaviruses SARS-CoV, and MERS-CoV which have high mortality rates. Our approach focuses on the identification of small molecule inhibitors that target conserved viral enzymes indispensable for viral replication, forestalling the evolution of resistance mutations.

Advantageous from a therapeutic perspective, the large +ssRNA genome of SARS-CoV-2 [1] also means the virus encodes a large number of proteins that may be potential drug targets: sixteen non-structural proteins (nsps), four structural proteins, and potentially nine accessory proteins (orfs) [2]. Primary searches for small molecule therapeutics have focused on the two main proteases encoded by SARS-CoV-2, nsp5 (Mpro) and nsp3 (PLpro) [3], and the major proteins of the RNA-dependent RNA-polymerase (RdRp) nsp12, and RdRp accessory proteins, nsp7 and nsp8 [2]. An overlooked target is nsp15, a protein associated with replication-transcription complexes that is conserved within the *Nidovirales* order with no corresponding host cellular counterpart.

Nidoviruses (*Coronaviridae, Arteriviridae*, and *Roniviridae*) are unique among RNA viruses in that they encode uridyl-specific endonuclease (endoU) [4]. The CoV nsp15 endoU RNase function has been demonstrated to be essential for efficient replication and transcription of viral proteins [5] and the evasion of double-stranded (ds) RNA viral recognition by host sensors in macrophages (reviewed in [6]). This is believed to be due to nsp15 cleaving viral dsRNA that ‘escapes’ virus-induced double-membrane vesicles (DMV) where normally replicating viral RNAs are sequestered. Inhibition of CoVs lacking nsp15 is manifest in macrophages or respiratory epithelial cells due to exuberant type I interferon (IFN) production, but not manifested in fibroblasts which do not activate IFN as readily. This is supported by studies showing that nsp15-deficient CoV activates multiple host dsRNA sensors independently, including innate immune pathways MDA5, PKR, and OAS (reviewed in [6]). A recent study reported that SARS-CoV endoU cleaves the variable lengths of 5’- polyuridines from negative-sense viral RNA (PUN RNA), which can otherwise act as a pathogen-associated molecular pattern (PAMP) by folding back and hybridizing with A/G rich domains, forming dsRNA-like secondary structures that can be recognized by host pattern recognition receptors like MDA5 [7]. Given that type 1 IFN is upregulated systemically during viral infection, it is expected that inhibition of nsp15 endoU RNase activity will lead to increased IFN sensitivity and diminished pathogenesis. Supporting this hypothesis, mouse hepatitis virus (MHV) (a related CoV) deficient in nsp15 endoU RNase function is highly attenuated in vivo compared with wild type virus [8]. However, some studies suggest SARS-CoV-2 infection is associated with less IFN production than other viral infections, so it is not clear how much interaction between IFN and nsp15 endoU RNAse will occur during SARS-CoV-2 infection [9, 10].

Seattle Structural Genomics Center for Infectious Disease (SSGCID), the Center for Structural Genomics for Infectious Diseases (CSGID), and at least two other groups have determined the structure of SARS-CoV-2 nsp15 (PDB IDs: 6XDH, 6VWW, 7KEH and 6K0R). SARS-CoV-2 nsp15 is structurally similar to SARS-CoV, MERS-CoV and MHV nsp15 structures [11–14]. This suggests that antivirals targeting SARS-CoV-2 nsp15 endoU may also similarly inhibit the highly conserved nsp15 endoU function in all beta-coronaviruses. Notably, no human homologs of this protein were identified when we conducted a DALI 3D-search, providing strong support that identifying selective inhibitors against nsp15 is a viable strategy. Although recent studies have identified the uracil derivative, tipiracil, as an inhibitor of SARS-CoV-2 nsp15 endoU activity, this compound exhibits only minimal antiviral activity at 50 μM concentrations, thus more potent inhibitors are desired [15]. Recently, a preprint chemically-genetically validated alpha-coronavirus nsp15 endoU as a target for antiviral activity [16]. Taken together, nsp15 endoU RNase appears to be an attractive therapeutic target for high-throughput screening (HTS) and structure-guided drug discovery (SGDD).

Given the challenges of developing new antivirals, identifying existing compounds that are already approved for use in the clinic or have progressed through early stages of clinical trials provides the opportunity to expedite SARS-CoV-2 treatments into the clinic. Therefore, to explore maximal repurposing opportunities for compounds that inhibit nsp15 function, we screened the ~13K ReFRAME library, assembled by Calibr/Scripps [17] and the Pandemic Response and COVID Box compound collections assembled by the Medicines for Malaria Venture (MMV) (mmv.org/mmv-open). The Calibr/Scripps ReFRAME repurposing library consists of both registered drugs (~40%) and clinical lead compounds that have progressed into Phase 1-3 clinical trials, as well as a smaller number of compounds from late pre-clinical development [17]. The MMV Pandemic Response Box library is a diverse set of 400 pharmacologically-active compounds against bacteria, fungi, and viruses, at various stages of drug discovery and development. The MMV COVID Box consists of 80 compounds (recently expanded to 160) that are demonstrated to have anti-SARS-CoV-2 effects. Thus, this collection of compounds represents an ideal screening set to find active compounds against SARS-CoV-2 nsp15.

## Results

### Nsp15 primary assay development

In order to identify small molecule inhibitors of the uracil-specific endoU function of nsp15, we adapted a previously described fluorescence resonance energy transfer (FRET) assay into a high-throughput format [12, 18]. Because nsp15 is reported to preferentially cleave uridylates (rU) [19], a four nucleotide chimeric substrate consisting of a single rU flanked by deoxyadenosines (dA) was utilized (**Fig 1A**). Nsp15 specific cleavage was monitored by appending a 5’ carboxyfluorescein (FAM) fluorophore and 3’ tetramethylrhodamine (TAMRA) quencher at opposing ends of the oligonucleotide. Enzymatically liberated 5’ FAM generates a fluorescence signal that can be detected on a fluorimeter. Moreover, this fluorescence-based system is amenable to 384-well format and 10 μL reaction volumes making it suitable for HTS of compound libraries.

**Fig 1.**
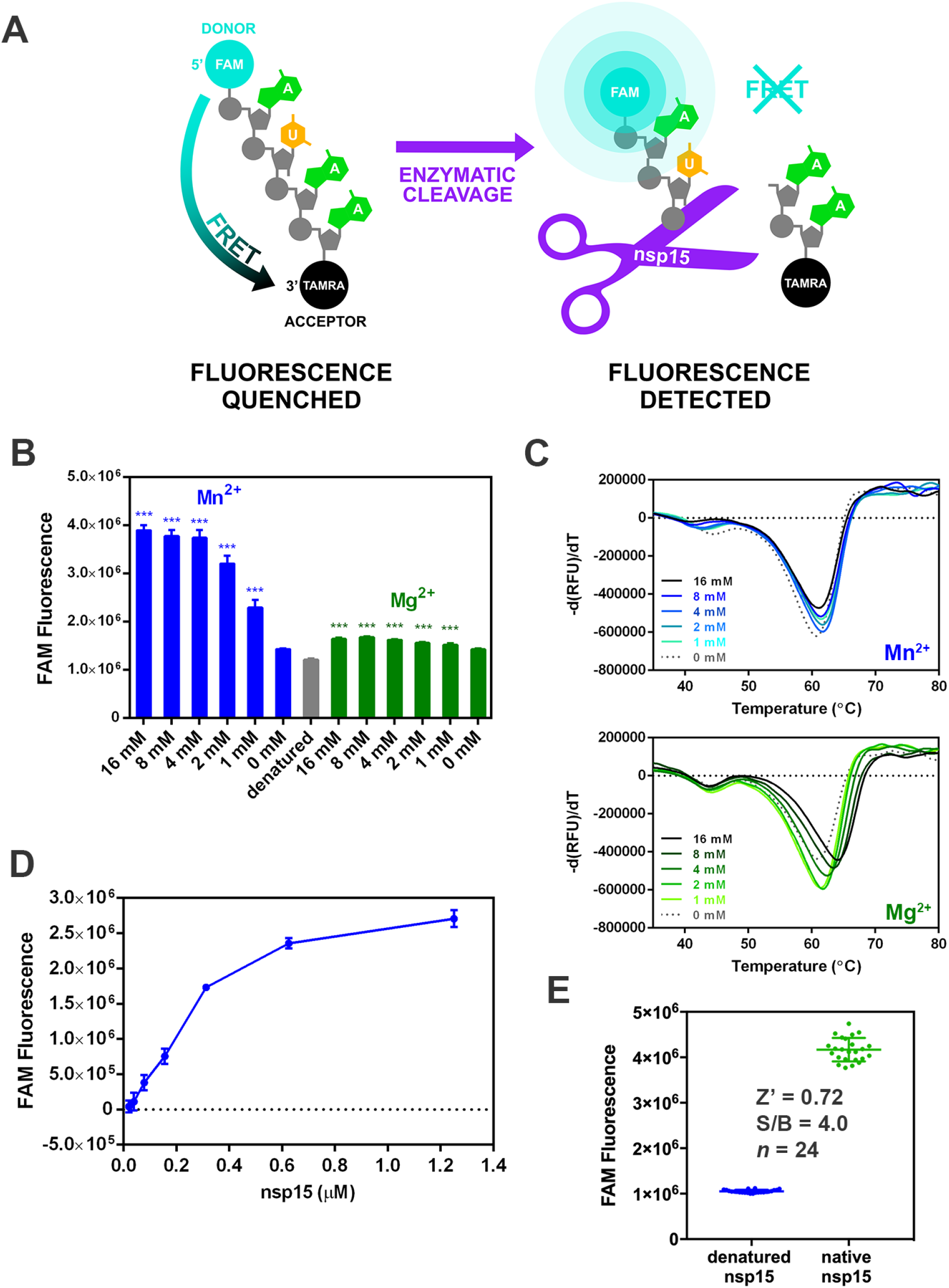
Primary SARS-CoV2 nsp15 endoribonuclease assay. **A)** Schematic illustrates the assay used to identify nsp15 inhibitors. Close proximity of the intact 5’-FAM and 3’-TAMRA fluorophores of the oligonucleotide substrate, 5’6-FAM/dArUdAdA/3’-TAMRA, results in quenching of fluorescence due to fluorescence resonance energy transfer (FRET). Enzymatic cleavage of the substrate by nsp15 leads to increased FAM fluorescence. **B)** The effects of Mn^2+^ and Mg^2+^ on SARS-CoV-2 nsp15 endoU activity in a 1 hr cleavage reaction at ambient temperature. The presence of Mn^2+^ leads to a 2.7-fold increase in FAM fluorescence compared to reactions lacking metal supplementation while the addition of Mg^2+^ appears to have minimal effect on activity (n=8, mean values ± standard deviations). ***, *P* < 0.001 (unpaired Student’s *t* test). **C)** Differential scanning fluorimetry (DSF) profiles of nsp15 in the presence of Mn^2+^ (blue) and Mg^2+^ (green). Graphs plotted as derivative of the rate of change of fluorescence (-d(RFU)/dT) versus temperature (°C), with the peak minima corresponding to the melting temperatures (T_m_). **D)** Thirty min cleavage reaction at ambient temperature using 1 μM 5’6-FAM/dArUdAdA/3’-TAMRA substrate and increasing nsp15 concentrations. **E)** FRET assay measuring nsp15 endoU activity under optimized assay conditions using 25 nM nsp15 and 0.5 μM 5’6-FAM/dArUdAdA/3’-TAMRA. The reaction was incubated for a 1 h at ambient temperature and fluorescence measured at excitation/emission wavelengths of 492 nm/518 nm on a microplate fluorimeter. Calculated Z’ and signal-to-background ratio (S/B) values are presented.

The optimal buffer conditions for Nsp15 endoU function were determined using differential scanning fluorimetry (DSF). In this assay, protein unfolding induced by thermal denaturation is measured by monitoring changes in binding of the hydrophobic fluorescent dye (SYPRO orange) which preferentially binds hydrophobic residues that are exposed as the protein unfolds. The temperature of the transition midpoint, or melting temperature (T_m_) exhibited a concentration dependent increase in the presence of Mg^2+^ (~3°C at 16 mM), while no such effect was observed with Mn^2+^ (**Fig 1C**). In other CoVs such as SARS-CoV, MERS-CoV, and MHV, Mn^2+^ was reported to significantly stimulate nsp15 endoribonuclease activity [12, 19], but not Mg^2+^. In line with these previous observations, a concentration dependent increase in SARS-CoV2 nsp15 activity was observed in the presence of Mn^2+^, with a 2.7-fold increase in FAM fluorescence observed between the no metal control and the highest concentration tested (16 mM). In contrast, Mg^2+^ concentrations up to 16 mM only minimally stimulated endoU activity (**Fig 1B**). Based on this data, we supplemented the endoU reaction buffer with 5 mM of MnCl2 in subsequent FRET HTS assays (**Fig 1D**). The enzyme and substrate concentrations were optimized to allow for a suitable dynamic range (>3-fold signal-to-background ratio) in a 1 hr reaction at ambient temperature (**Fig 1E**). A commonly used statistic is the Z’ score, a measurement that is reflective of an assay’s dynamic range and data variation of neutral and baseline controls, with an “excellent” assay defined by a Z’ score between 0.5 and 1 [20]. The final 384-well assay, using 25 nM enzyme and 0.5 μM substrate, yielded a robust Z’-score of 0.72 from three optimized experiments.

### Nsp15 high-throughput screen of ReFRAME, Pandemic Response Box, and COVID Box libraries

A primary screen at an assay concentration of 10 μM was performed against Calibr/Scripps ReFRAME and the MMV Pandemic Response Box and COVID Box Libraries. The overall assay performance was robust, with average Z’-scores of 0.75 ± 0.08 for the ReFRAME library, 0.70 ± 0.06 for the Pandemic Box, and 0.61 for the COVID Box (**Fig 2**). Of the 13,161 compounds in the ReFRAME library, 23 compounds inhibited nsp15 cleavage activity by >50%, representing a hit rate of 0.17%. Of the 400 compounds in the Pandemic Response Box, one compound inhibited nsp15 activity by >50%, corresponding to a hit rate of 0.25%. No hits were observed in the 80 compounds of the COVID Box.

**Fig 2.**
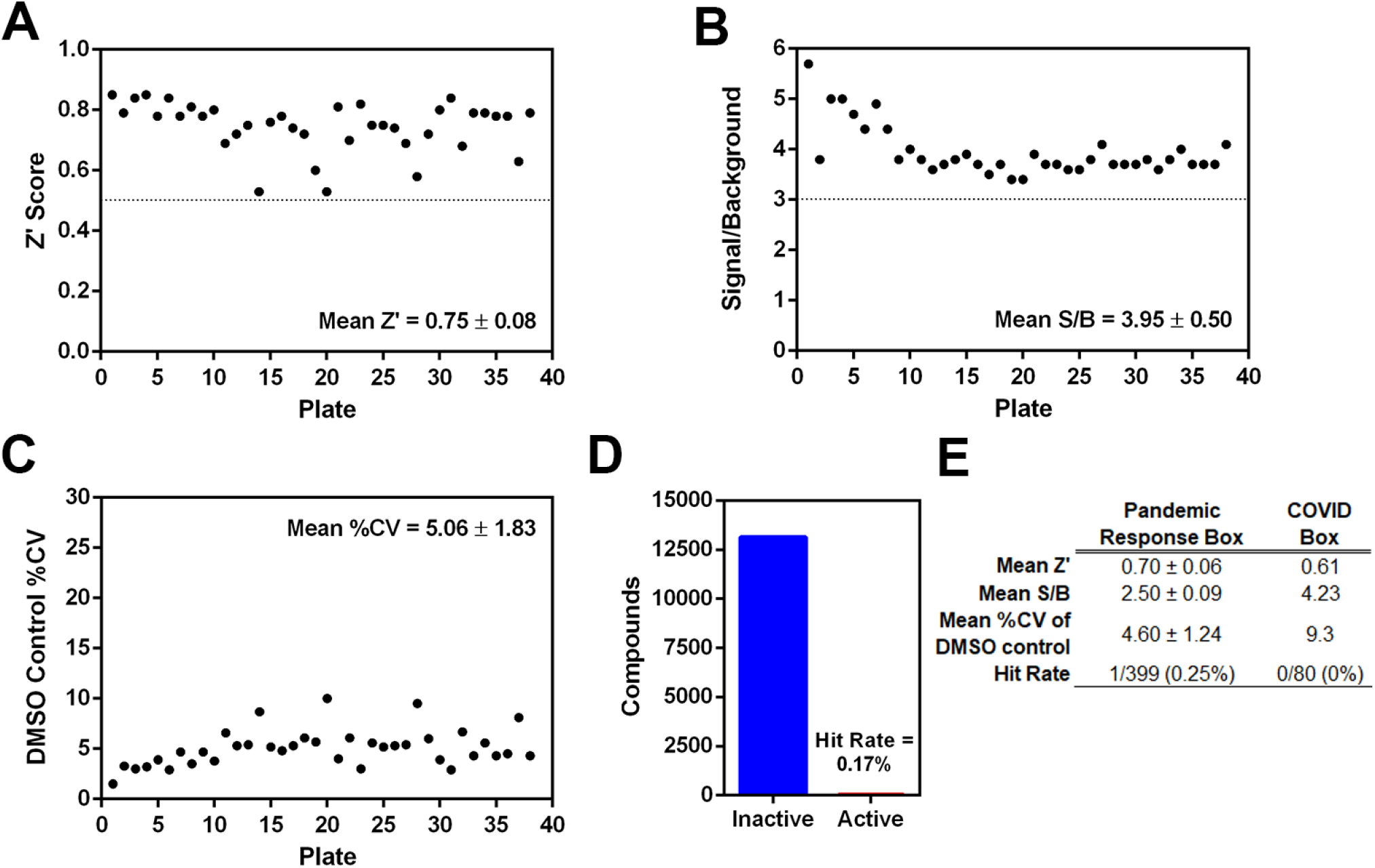
Results of a primary high-throughput screen against the ReFRAME, Pandemic Response and COVID Box libraries. **A-D)** Assay statistics for the ReFRAME library plotted for **A)** mean Z’ score, **B)** mean signal-to-background (S/B) ratio, **C)** mean %CV of DMSO controls, and **D)** overall hit rate. **E)** Summary of the mean Z’ scores, S/B ratios, and %CV values for the Pandemic Response Box and COVID Box libraries.

All hits were validated using the same assay conditions with compounds prepared as 8-point, 3-fold dilution series (pre-spotted onto 384-well plates by Calibr) and a top concentration of 10 μM for confirmation in a secondary dose response assay. Out of 23 primary hits from the ReFRAME library, 15 were confirmed (62.5%) as recapitulating the activities of the 10 μM screen (**Table 1**). Two of these hits were identified as the tissue staining agent Trypan Blue and were duplicate entries in the ReFRAME collection, eliminating them from further consideration. The remaining 13 molecules were sourced from Calibr or commercial vendors and subjected to a tertiary FRET assay at higher concentrations to determine half-maximal inhibitory concentration (IC_50_) values. Compound PF-06260414 failed to exhibit >50% activity at concentrations up to 80 μM and was therefore deprioritized. Ceftazidime, which was sourced as a hydrated salt and dissolved in water, showed reduced potency in the tertiary assay with an IC_50_ of 72 μM and was also deprioritized. Compound BVT-948 exhibited the highest potency with an IC_50_ of 0.1 μM. The tertiary assay identified several compounds with submicromolar IC_50s_ (Streptonigrin, β-lapachone, and Tesimide) and compounds with IC_50s_ <10 μM (Exebryl-1, BN-82685, Azaquinone, Piroxantrone, Pirarubicin, and Doxorubicin), as summarized in **Table 1**. The sole Pandemic Response box hit, MMV1580853, was sourced from MMV and reconfirmed with an IC_50_ of 11.5 μM.

**Table 1.** Biochemical assay results and attrition of hits.

|  | 1° FRET Assay<br>% inhibition | 2° FRET Assay<br>IC<sub>50</sub> (μM) | 3° FRET Assay<br>IC<sub>50</sub> (μM) | DSF Assay<br>ΔT<sub>m</sub> (°C) | Amplex Red Assay<br>% Activity | RNA Cleavage Assay<br>IC<sub>50</sub> (μM) |
| --- | --- | --- | --- | --- | --- | --- |
| **Exebryl-1** | 57 | 1.06 | 1.28 | 0.1 | 0 | 9.27 |
| **Piroxantrone** | 74 | 3.95 | 2.46 | -1.1 | 0 | 9.1 |
| **MMV1580853** | 62 | >10 | 11.5 | 0.1 | 21 | NA |
| **pirarubicin** | 66 | 3.78 | 5.98 | 0.4 | 51 |  |
| **Doxorubicin** | 57 | 7.46 | 9.29 | -0.5 | 52 |  |
| **BVT-948** | 90 | 4.96 | 0.1 | -11.8 | >100 |  |
| **Streptonigrin** | 89 | 4.25 | 0.29 | -11.5 | >100 |  |
| **β-Lapachone** | 73 | 2.78 | 0.31 | -11.5 | >100 |  |
| **Tesimide** | 82 | 1.09 | 0.75 | -11.7 | >100 |  |
| **BN-82685** | 76 | 6.32 | 1.43 | -11.3 | >100 |  |
| **Azaquinone** | 80 | 3.12 | 1.99 | -11.4 | >100 |  |
| **Mecobalamin** | 58 | 3.34 | 18.3 | -3.4 | >100 |  |
| **Ceftazidime** | 66 | >10 | 72 |  |  |  |
| **Trypan Blue** | 58 | 5.08 |  |  |  |  |
| **PF-06260414** | 74 | 7.69 |  |  |  |  |
| **Trypan Blue** | 51 | 7.78 |  |  |  |  |
| **L 658310** | 54 |  |  |  |  |  |
| **Ganstigmine** | 51 |  |  |  |  |  |
| **Carubicin** | 56 |  |  |  |  |  |
| **Aminoquinuride** | 95 |  |  |  |  |  |
| **mitoxantrone** | 73 |  |  |  |  |  |
| **vecuronium bromide** | 61 |  |  |  |  |  |
| **Sepantronium bromide** | 50 |  |  |  |  |  |

Hits from the initial 10 μM primary screen were subsequently tested in a secondary dose response assay with a top concentration of 10 μM. Active hits were then reconfirmed in a tertiary dose-response assay using an expanded concentration range. Differential scanning fluorimetry and Amplex Red assays further eliminated compounds with undesirable redox activity. Finally, 3 compounds were tested in an orthogonal cleavage assay, and two compounds, Exebryl-1 and Piroxantrone, demonstrated inhibition of endoU activity. All values are derived from one experiment except for the RNA cleavage assay, which was performed in *n*=3 technical replicate experiments.

Abbreviation: FRET, fluorescence resonance energy transfer; DSF, differential scanning fluorimetry; IC_50_, half maximal inhibitory concentration; ∆T_m_, shift in melting temperature; NA, not available due to poor compound solubility.

### Nsp15 hit characterization, reactive compounds, and orthogonal assays

Binding interactions of hit compounds with nsp15 was examined with a DSF assay. In the assay, interactions of small-molecule ligands with target proteins will often result in improved thermal stability, which is reflected by an increase in protein T_m_. None of the inhibitors produced a significant positive shift (>2°C) in T_m_. Instead, several inhibitors were observed to produce a significant negative T_m_ shift, which is suggestive of a destabilizing effect (**Table 1**). Streptonigrin, μ-lapachone, BVT-948, Azaquinone, BN-82685, and Tesimide all lowered the nsp15 Tm by more than 10°C. The remaining inhibitors did not exhibit significant shifts in the nsp15 T_m_. Among the inhibitors that produced a strong negative T_m_ shift, many were observed to be quinone-containing compounds, which are commonly regarded as pan-assay interference compounds (PAINS). Quinone-containing compounds readily participate in redox-cycling reactions in the presence of thiol-based reducing agents and generate reactive oxygen species (ROS) such as H_2_O_2_, resulting in non-specific protein inhibition [21].

Because the FRET and DSF assay buffers contained the reducing agent, dithiothreitol (DTT), the prospect for redox cycling was deemed high for these inhibitors. Therefore, the Amplex Red assay was employed as a counter screen to measure the generation of H_2_O_2_ in the presence of 1 mM DTT (**Fig 3**). The Amplex Red reagent reacts with H_2_O_2_ in a 1:1 stoichiometry to produce an oxidized, red-fluorescent product, resorufin, which can be detected on a fluorimeter at excitation/emission wavelengths of 531/595 nm. All compounds that elicited a strong negative T_m_ shift in the DSF assay were observed to be active in the Amplex Red assay, suggesting these hits were thiol-reactive compounds that may be contributing to non-specific inhibition of nsp15 endoU enzyme activity in the FRET assay and the observed destabilization of nsp15 in the DSF assay (**Table 1**). In addition to the quinone-containing compounds, BVT-948, Mecobalamin and Tesimide were also identified as redox reactive compounds. These reactive compounds were deprioritized for further studies. Piroxantrone and Exebryl-1 did not exhibit significant redox activity, while MMV1580853 retained minimal activity (<25%). We chose to prioritize these compounds for further investigation.

**Fig 3.**
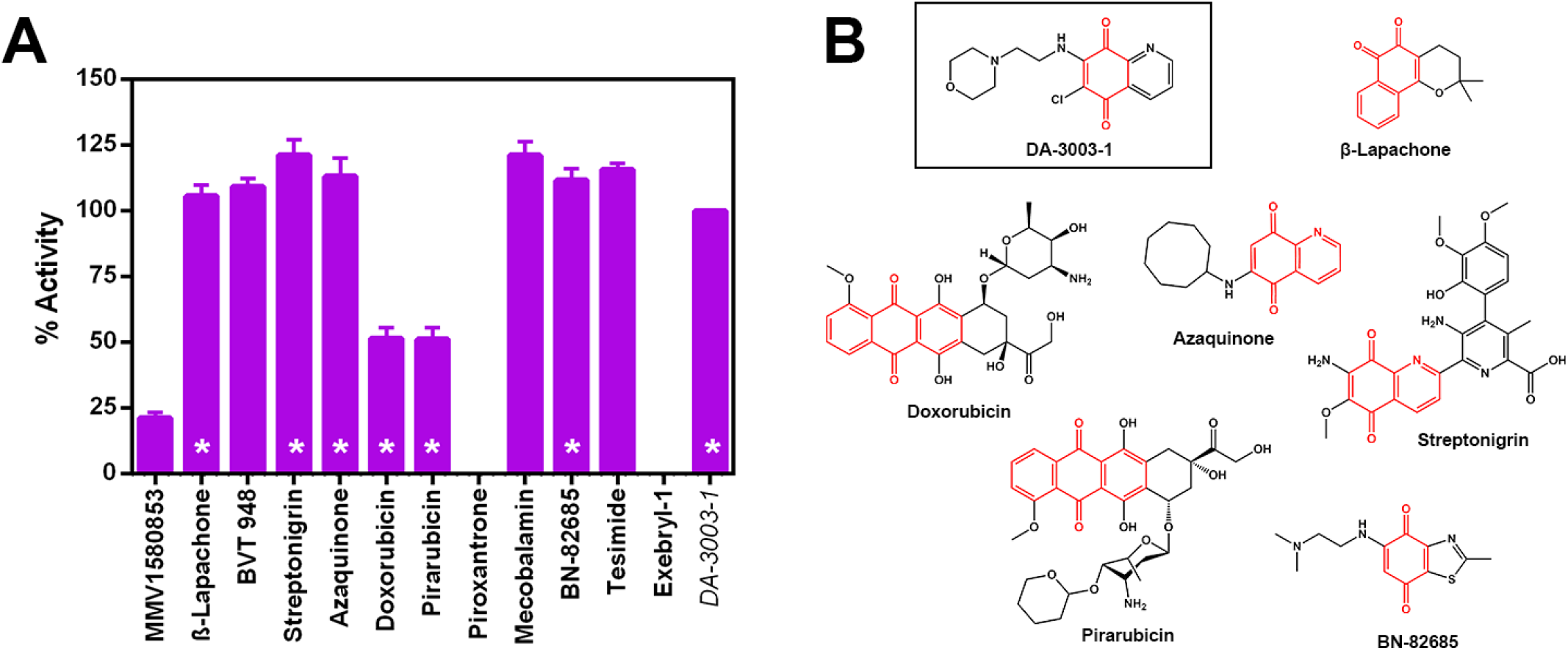
Amplex Red counter screen for potential redox activity of select hit molecules. **A)** Plot of the redox cycling activity of hit compounds using the Amplex Red assay in the presence of 1 mM DTT when compared with a known redox-cycling reference compound, DA-3003-1. Asterisks denote compounds that contain a quinone group. **B)** Structures of quinone-containing compounds with the quinone group highlighted in red.

Additional validation of Piroxantrone, Exebryl-1 and MMV1580853 inhibition of nsp15 endoribonuclease function was conducted using a non-fluorescence, gel-based RNA cleavage assay (**Fig 4**). Specifically, a 31 nt poly(rA) RNA oligonucleotide substrate with a single rU cleavage site was utilized, which upon cleavage by nsp15 is converted to 20 nt and 10 nt products. The cleavage assay was typically run by incubating the RNA oligonucleotide with nsp15 and varying concentrations of inhibitor for 2 h at ambient temperature prior to analysis of the reaction products on polyacrylamide gels and staining with SYBR green II to visualize the resultant RNA bands. EndoU activity in the presence of inhibitor was quantified by measuring the relative abundance of cleaved and uncleaved poly(rA) relative to nsp15 treated with DMSO. Exebryl-1 and Piroxantrone demonstrated concentration-dependent inhibition of cleavage, congruent with the results of the FRET assay. However, MMV1580853 exhibited poor solubility and readily precipitated out of solution when diluted in assay buffer, therefore, its activity in the RNA cleavage assay was not replicable. The IC_50_ values calculated from the RNA cleavage assay (9.27 μM for Exebryl-1 and 9.10 μM for Piroxantrone) were consistent with calculated IC_50_ values from the FRET assay (**Fig 4A-D**, **Table 1**).

**Fig 4.**
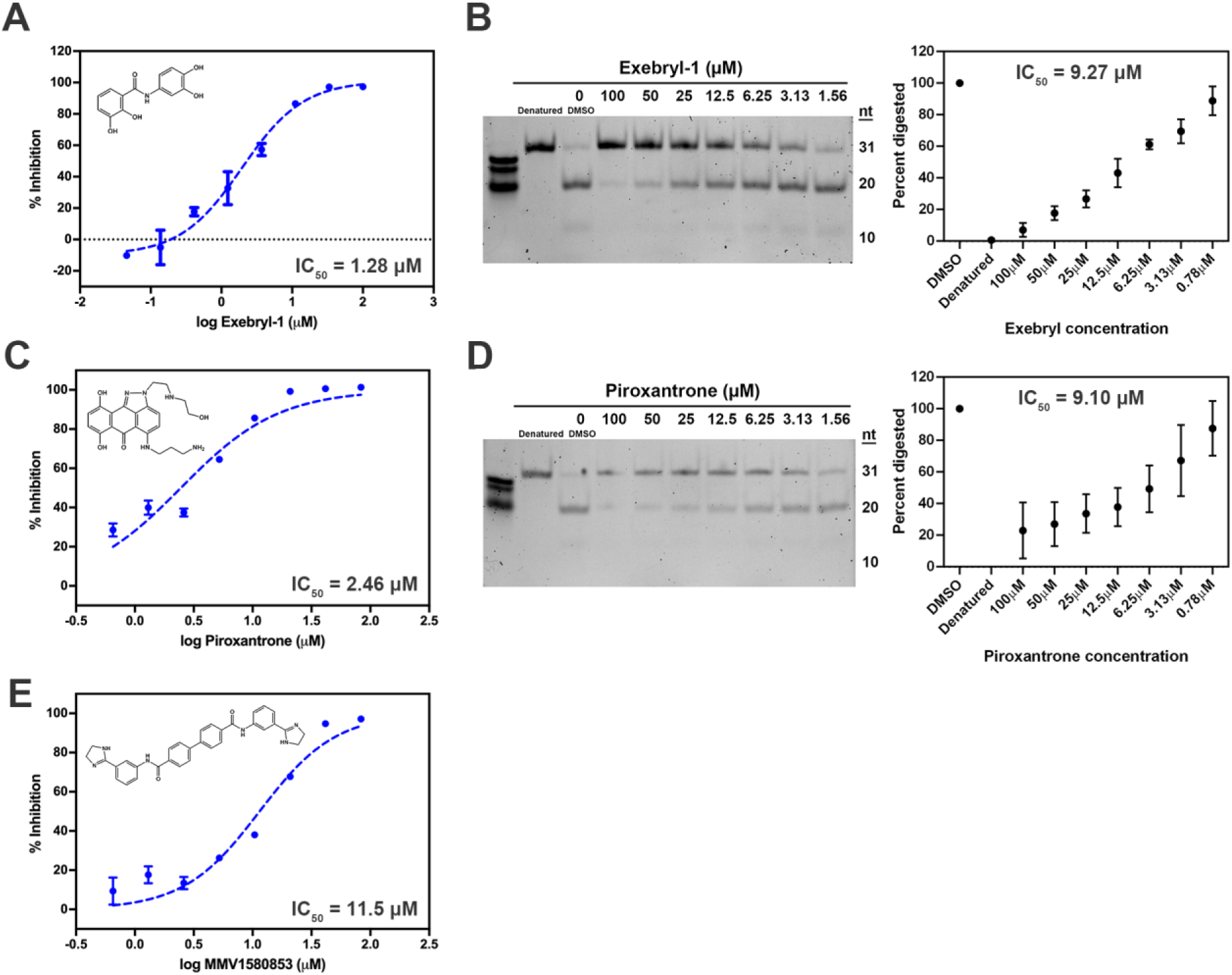
Dose response curves of select nsp15 hits. Inhibitor structures and 3° FRET assay dose-response curves with calculated IC_50_ values for Exebryl-1 (**A**), Piroxantrone (**C**), and MMV1580853 (**E**). Nsp15 (25 nM) was incubated with 2-fold or 3-fold serial concentrations of compound and incubated with the substrate 5’6- FAM/dArUdAdA/3’-TAMRA (0.5 μM) at ambient temperature for 1 h. Results of a polyacrylamide gel-based RNA cleavage assay for Exebryl-1 (**B**) and Piroxantrone (**D**). nsp15 (0.78 μM) and poly(rA) (500 ng) was incubated with 2-fold serial concentrations of each compound and incubated at ambient temperature for 2 h. Cleavage products were separated from uncleaved poly(rA) by electrophoresis on a 15% TBE-urea polyacrylamide gel and the % cleavage activity calculated by quantitating the relative amount of cleaved (20 nt lower band) vs uncleaved (31 nt top band) substrate in each sample relative to a DMSO control. Gel-based RNA cleavage inhibition of MMV1580853 was observed, but was variable from experiment to experiment, likely due to poor solubility in DMSO and assay buffer.

### Native Mass Spectrometry (MS) and molecular docking of nsp15 hits

As the negative T_m_ shifts observed in the DSF binding studies suggested that binding of compound might lead to destabilization of nsp15, we used native mass spectrometry (MS) to further investigate binding activities for select inhibitors. Native MS has successfully been applied to confirm inhibitor binding for the main SARS-CoV-2 protease Mpro [22–24]. In native MS, nsp15 was electrosprayed from non-denaturing ammonium acetate solution in the presence of an inhibitor. Inhibitor binding can be evaluated by monitoring the increase in mass of the inhibitor-nsp15 complex compared to the apo (protein with no ligand) nsp15 protein. The native nsp15 hexamer exhibited an observed mass of 241.8 kDa (**Fig 5A**). The high mass and heterogeneity of nsp15 increases the difficulty of distinguishing inhibitor binding, especially on lower resolution mass spectrometers. Therefore, we performed ion activation on mass-isolated nsp15 hexamers to release smaller protein subunits allowing us to better resolve inhibitor binding. Collision-induced dissociation (CID), widely available in commercial mass spectrometers, involves acceleration of protein ions into a pressurized collision cell. Upon CID, the internal energy of the protein increases from multiple collisions with neutral gas molecules. However, this “slow heating” process typically induces protein unfolding, release of subunits, and (partial) loss of weakly bound ligands [25]. To better capture the inhibitor binding profile, we also applied surface-induced dissociation (SID), which involves collisions with a surface. Distinct from CID, SID is considered a “fast heating” process, which generally induces less unfolding and allows for better ligand preservation on released subunits [25, 26].

**Fig 5.**
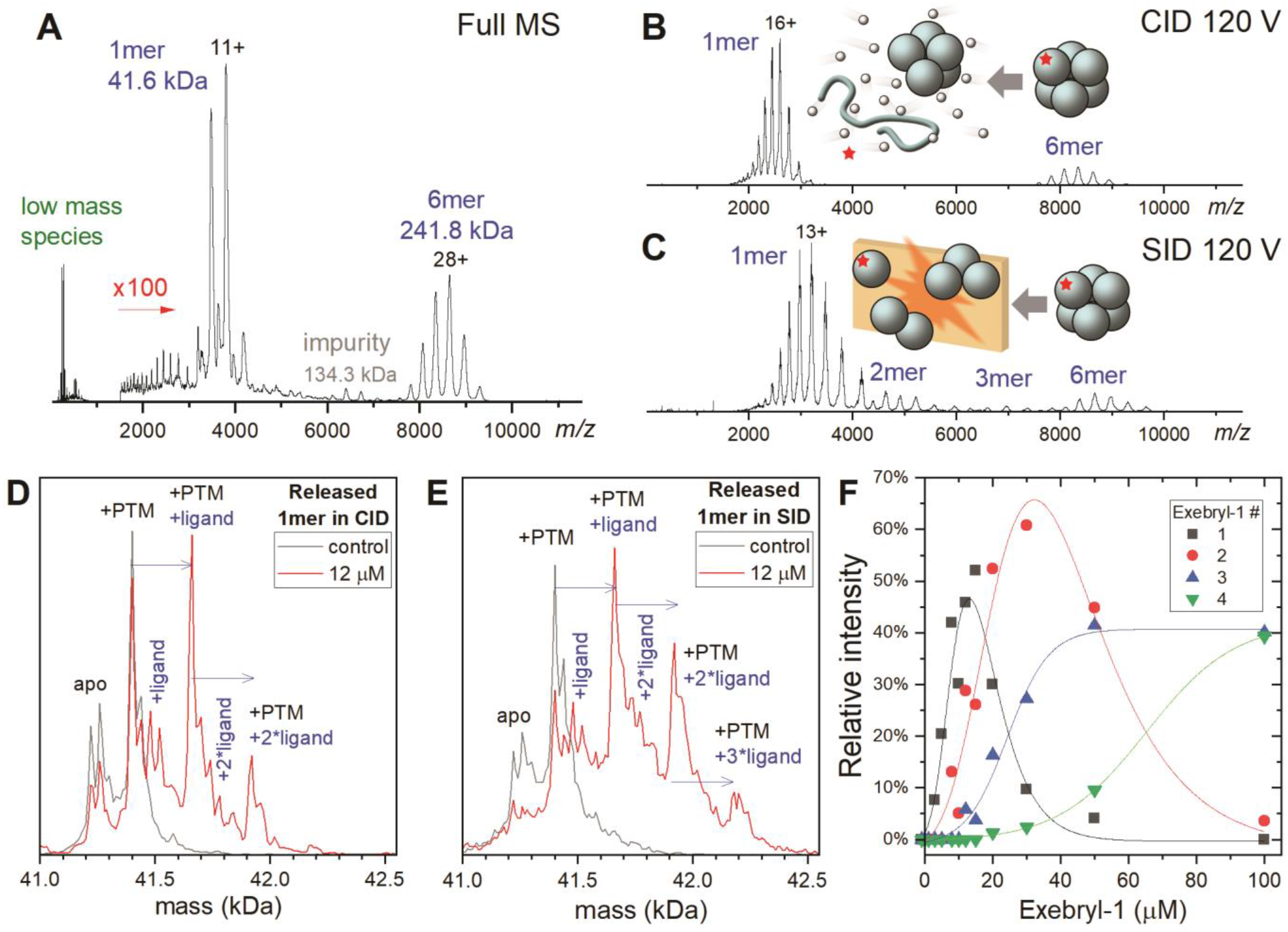
Native MS of nsp15 and its binding to Exebryl-1. **A)** Representative native MS spectrum of nsp15. Low mass species such as residual buffer molecules (and excess ligand if any) are detected below *m/z* 1000. Nsp15 hexamers (6mer) and monomers (1mer) were seen around *m/z* 8000-9000 and 2000-4000, respectively. Spectrum > *m/z* 1500 is magnified 100 times for clarity. **B)** Collision induced dissociation (CID) and **C)** surface induced dissociation (SID) of isolated high mass 6mers released 1mers. SID also released dimers (2mer) and trimers (3mer). Other low intensity species are not labeled. The insert cartoon illustrates the different dissociation pathways accessed by CID (slow heating) and SID (fast heating). SID typically induces less unfolding and loss of weakly bound ligands. **D-E)** Deconvoluted mass distribution of the released 1mers by CID and SID, respectively. The control sample (gray trace) showed the apo 1mer and another peak from post-translational modification (+PTM) of ~178 Da. With 12 μM Exebryl-1 (red trace), both CID and SID showed new peaks compared to the control which were assigned to 1-3 Exebry-1 binding events (mass shift of 261 Da per binding). **F)** Relative intensity of SID-released nsp15 1mers with different numbers of Exebryl-1 molecules as a function of the Exebryl-1 concentration.

By increasing the acceleration voltage to 120V, CID and SID of the nsp15 hexamer both released primarily monomeric species (**Fig 5B-C**), with SID resulting in lower charged monomers and additional dimers/trimers. By examining the deconvoluted mass profiles of the released monomers, Exebryl-1 binding could be detected as shown by the representative data in **Fig 5D-E**. The control samples (gray traces) showed two major peak clusters. The peak cluster centered at 41.2 kDa is the apo protein (sequence mass and salt adducts). The second cluster at 41.4 kDa is the apo protein with post-translational modification (PTM) of ~178 Da, tentatively assigned to an N-terminal gluconoylation species, a common artifact on recombinant proteins expressed in *Escherichia coli* [27]. At 12 μM Exebryl-1, both CID- and SID-released monomers showed additional peaks with mass shifts corresponding to 1-3 bound ligands (Exebryl-1, 261 Da). As expected, SID- released monomers retained more bound ligands. **Fig 5F** shows the relative intensity of the SID-released monomers with varying number of Exebryl-1 molecules bound. The *K_d_* for the first binding event was estimated to be ~12 μM, where the apo and single Exebryl-1 bound monomers species are at similar intensities. Interestingly, Exebryl-1 showed continuous binding to nsp15 as its concentration was increased. At 100 μM, five and six molecules of Exebryl-1 were detected at ~10% each (not plotted in **Fig 5F**). The high and heterogeneous binding stoichiometry imply multiple binding sites.

Another possibility we considered is that Exebryl-1 binding to nsp15 is asymmetric and concentrated on one or a few monomers, which are preferentially released by CID and SID from the nsp15 hexamer. We thus more thoroughly examined the mass addition on the nsp15 hexamers in the SID spectra (**Fig S1**). Assuming the mass addition was exclusively from Exebryl-1, the number of Exebryl-1 molecules bound to the hexamer matched well with six times the average bound Exebryl-1 in the released monomer. These result suggests the binding was symmetric among the six monomers in the complex. The same trend was observed for the CID data, but the average number of ligands were consistently lower than those observed from SID. This difference can be attributed to the higher degree of unfolding and loss of weakly associated ligands in CID compared to SID. We also performed denaturing LCMS and peptide mapping (trypsin digestion) of nsp15 incubated with Exebryl-1. No significant Exebryl-1 modified protein or peptides were observed, suggesting binding is likely non-covalent.

Using the same CID and SID analyses as conducted for Exebryl-1, Piroxantrone and MMV1580853 were also examined. Unlike Exebryl-1, these two compounds exhibited binding affinities that were too weak to be detected on CID- or SID-released monomers at ~ 10 μM. It is likely most of the compounds were directly stripped off from the nsp15 hexamer upon activation, and the protonated compounds were observed among the low mass species. Representative data for Piroxantrone and MMV1580853, shown in **Fig S2**, indicate their weaker non-covalent binding to the nsp15 hexamer.

Attempts to obtain co-crystal structures of nsp15 bound to Exebryl-1, Piroxantrone, and MMV1580853 were unsuccessful. However, the availability of multiple high resolution X-ray crystal structures of SARS-Cov-2 nsp15 allowed us to predict the possible binding sites of Exebryl-1, MMV1580853, and Piroxantrone on nsp15. The compounds were docked to the SARS-CoV-2 nsp15 X-ray structure (PDB ID: 6XDH) using an automated Qvina docking workflow [28]. As a first step, the nsp15 X-ray structure was prepared for the calculations by removing all non-protein atoms in the crystal structure. Next, a pocket-selection tool (Qvina-W for blind docking [29]) was used to identify the most likely ligand binding pockets in the C-terminal catalytic domain surrounded by His235 and His250 in the nsp15 monomer. This approach is consistent with previous studies where a similar region was used for the nsp15 docking calculations [16]. Using Exebryl-1 and the identified pockets, docking scores were generated.

Two potential binding sites for Exebryl-1 were identified. In the higher scoring binding mode, Exebryl-1 is buried deep within a pocket formed between the C-terminal catalytic domain and N-terminal oligomerization domain, forming multiple hydrogen bonds with the residues Lys71, Thr275, and Tyr279 (**Fig 6A-C**). In the catalytically active hexameric form in which monomers are assembled into a dimer of trimers, this pocket is oriented within a central pore formed by the dual trimer rings (**Fig 6D**). In the second ranked binding mode (relatively less stable by only ~0.8 kcal/mol), Exebryl-1 is bound within the shallow groove of the endoU active site, forming hydrogen bonds with the side chains of the catalytic triad consisting of His235, His250, and Lys290, and with the backbone of Thr341 and Tyr343 (**Fig 6E-G**). The small energy differences between the two binding modes suggest that Exebryl-1 may bind to multiple sites on nsp15, which is supported by the native MS data. While the docking studies showed Exebryl-1 bound deep into multiple potential binding pockets of nsp15, similar calculations showed that MMV1580853 and Piroxantrone bound to nsp15 but did not bury themselves deep within the pockets (**Fig S3**).

**Fig 6.**
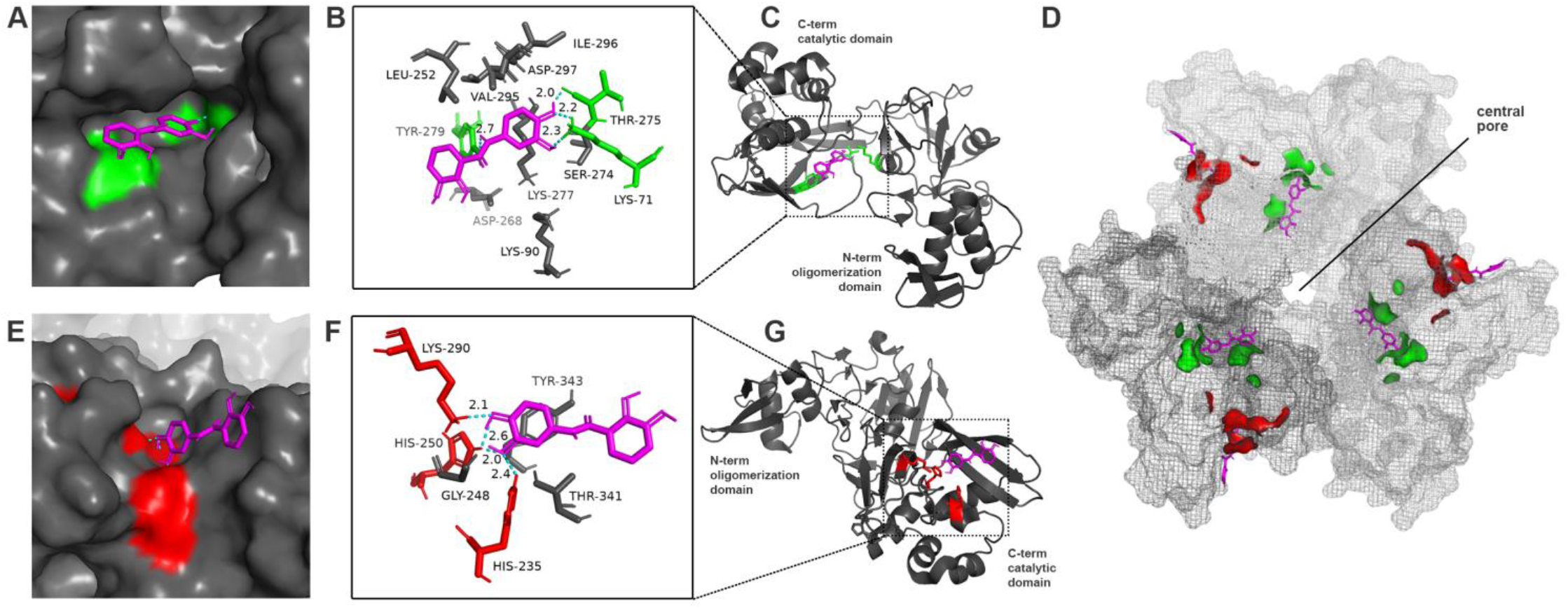
Predicted binding modes of Exebryl-1 to SARS-CoV-2 nsp15. Nsp15 is depicted in gray and Exebryl-1 in magenta. **A-C)** The top scoring binding model shows Exebryl-1 buried deep in a pocket formed between the C-terminal endoU and N-terminal oligomerization domain, forming hydrogen bonds with Lys71, Thr275, and Tyr279. Key residues indicated in green. Dashes indicate hydrogen bonds. **D)** In the trimer assembly, the pocket (green) is oriented within the central pore distant from the active site catalytic triad (red). Exebryl-1 binding to this pocket does not occlude the pore but rather faces towards it. **E-G)** The second ranking binding model (~0.8 kcal/mol less stable) shows Exebryl-1 bound to the endoU active site, forming hydrogen bonds with the catalytic triad His235, His250, and Lys290 (red), and the backbone of Thr341 and Tyr343.

### Cytotoxicity and antiviral activities of nsp15 hits

Exebryl-1, Piroxantrone, and MMV1580853 were assessed for safety against CRL-8155 (human lymphocyte) and HepG2 (human hepatocyte) cell lines. Exebryl-1 exhibited a CC_50_ of 37.8 μM in CRL-8155, but was safe up to 200 μM in HepG2 cells. Piroxantrone exhibited significant cytotoxicity, with a CC_50_ of <1.25 μM in CRL-8155 cells and 33.9 μM in HepG2 cells. MMV1580853 did not exhibit cytotoxicity (>80 μM) in either CRL-8155 or HepG2 cells.

Exebryl-1, Piroxantrone, and MMV1580853 were investigated in a primary cytopathic effect (CPE) assay in Vero kidney epithelial cells to screen for protection against SARS-CoV2 viral induced cell toxicity. When CPE was analyzed via microscopic observation, Exebryl-1 exhibited a modest EC50 (concentration that reduces CPE by 50%) of 10 μM and a CC_50_ (concentration that causes cytotoxicity, or reduced growth, by 50%) of 52 μM on Vero cells, with a selectivity index (SI) of 5.2. When CPE was assessed with neutral red staining in the same experiment with Exebryl-1, the EC50 was 16 μM and the CC_50_ of VERO cell growth was 57 μM, with a selectivity index (SI) of 3.6. Neither Piroxantrone nor MMV1580853 exhibited protection against viral CPE in this assay, although detection of viral CPE was limited by the low CC_50_ for Piroxantrone and low solubility for MMV1580853. Exebryl-1 was subsequently tested in a secondary viral yield reduction (VYR) assay to evaluate its ability to inhibit virus production in Caco-2 human colorectal adenocarcinoma cells.

Exebryl-1 demonstrated antiviral effects with an EC90 (concentration that reduces viral replication by 90%) of 51 μM via microscopic observation and CC_50_ of 61 μM in Caco-2 cells via neutral red staining, with a SI of 1.2. Next, Exebryl-1 was investigated for SARS-CoV-2 antiviral activity in Calu-3 human lung epithelial cells. A dose-dependent reduction in infection was observed via immunostaining, exhibiting an EC50 of 65.6 μM and CC_50_ of >200 μM, representing a SI of >3 (**Fig 7**).

**Fig 7.**
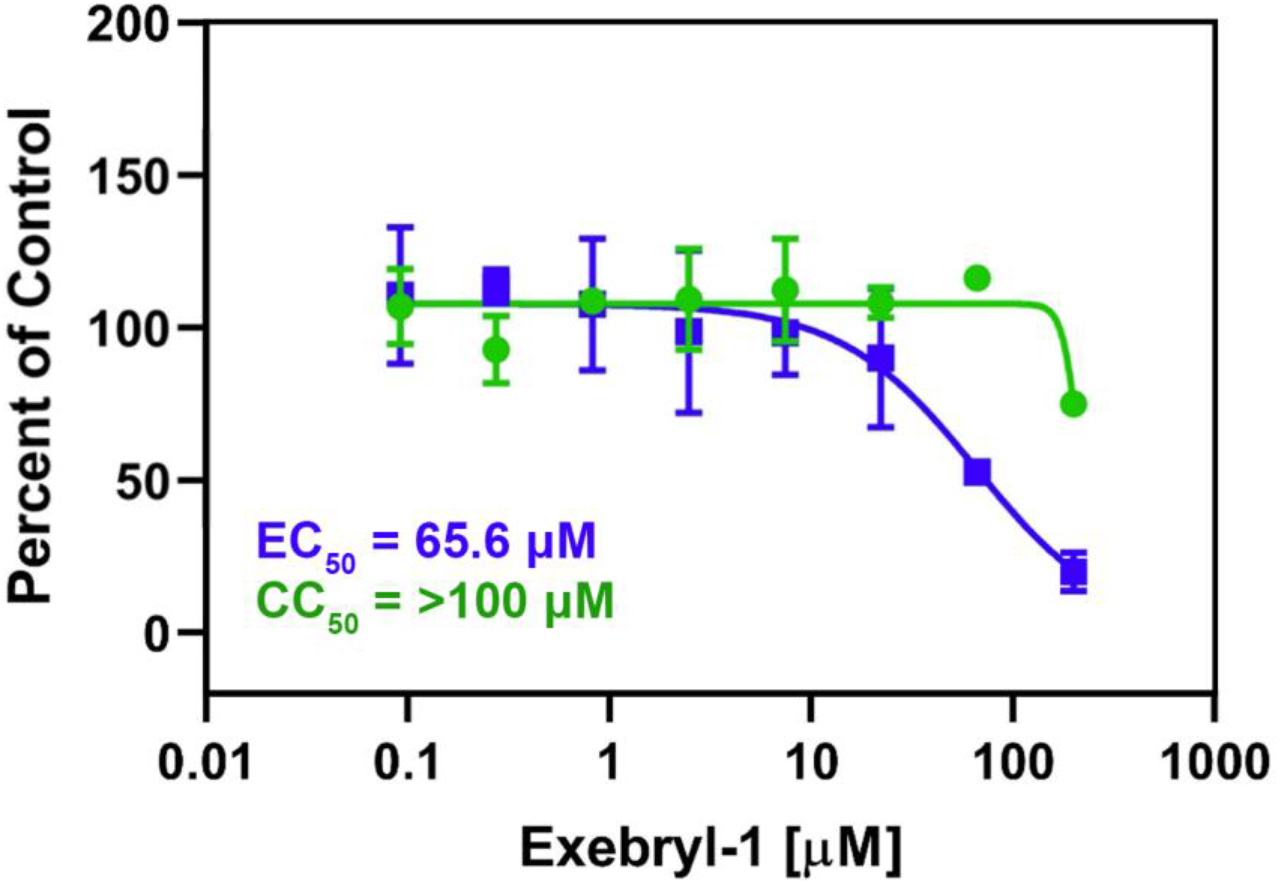
In vitro antiviral SARS-CoV-2 assay in Calu3 cells. Calu3 cells were pretreated with Exebryl-1 for 2 then infected with SARS-CoV-2 at a MOI=0.5. After 48 h, the cells were fixed, immunostained, and imaged by automated microscopy. Data were normalized to DMSO controls to determine EC50 (blue) and CC_50_ (green). Values represent average percentage of control for n=3 replicates.

## Discussion

Expressed and purified recombinant SARS-CoV-2 nsp15 was demonstrated to possess an uracil-specific endoU function using both a FRET and RNA in vitro assay. The FRET assay (1°) was developed into a HTS that was used to initially identify 23, 1, and 0 nsp15 inhibitors from the ReFRAME, Pandemic Response Box, and COVID Box libraries, respectively, using a cut-off threshold of 50% inhibition for hits to reduce the selection to inhibitors with IC_50s_ below 10 μM. After hits were reconfirmed in two subsequent FRET assays, this list was reduced to 12 compounds (11 ReFRAME plus 1 Pandemic Response Box). To verify ligand binding to nsp15, these 12 compounds were then analyzed with a DSF assay under the hypothesis that bound ligands increased the T_m_ of nsp15. Nine of the 12 compounds tested showed a decrease in the nsp15 T_m_ and none showed a significant positive T_m_ shift (>2°C). The observation that many of the compound resulted in a negative T_m_ shift (six > 10°C) and the realization that many of these hits were quinones lead us to analyze these compounds with an Amplex Red assay to detect H_2_O_2_ evolution. This is because quinones are often PAINs, false-positive molecules that generate ROS H_2_O_2_ which is responsible for protein destabilization. Such an analysis resulted in the elimination of nine of the 12 hits from further study leaving just three compounds for consideration: Exebryl-1, Piroxantrone, and MMV1580853. Inhibition of nsp15 endoU activity was confirmed for Exebryl-1 and Piroxantrone with an in vitro, non-fluorescent RNA cleavage assay using a single-stranded 31 nt poly(rA) containing a single rU. For MMV1580853, the RNA assay was non-replicable likely due to poor solubility in DMSO and assay buffer. While at least two of the final three hits showed promise towards inhibiting nsp15, the DSF assay failed to verify ligand binding for any of these compounds. Although we expect ligand binding to stabilize nsp15, the high apparent T_m_ (~61°C) for the 242 kDa hexamer may make additional stabilization by a small, 261 Da ligand difficult to detect. Therefore, ligand binding was probed using high resolution native MS where nsp15 was observed to bind Exebryl-1 with a *Kd* of ~12 μM for the first binding event (per monomer). Additional binding proceeds up to an average of approximately four ligands per monomer at a 100 μM Exebryl-1 concentration. Association of Piroxantrone and MMV1580853 to nsp15 was detectable by native MS, but the measured binding was significantly weaker relative to Exebryl-1 (**Fig S2**). Attempts at co-crystallography with each of the final three hits failed to identify crystals that had a density consistent with ligand binding. In silico molecular docking experiments with the three final hits and a crystal structure of nsp15 (PDB ID: 6XDH) showed that only Exebryl-1 bound deeply into potential binding pockets near the endoU active site and at multiple regions with similar affinities. The latter observations were consistent with the native MS experiments demonstrating binding of multiple Exebryl-1 molecules within each nsp15 monomer. The top scoring binding pose was identified to be in a deep pocket oriented within a central pore structure formed by the catalytically active hexamer, and a slightly less stable second binding pose was identified within the endoU catalytic site, forming hydrogen bond interactions with the catalytic triad. Although Piroxantrone and MMV1580853 were also observed to bind in the same sites, both compounds appeared to bind near the surface rather than buried within the pockets like Exebryl-1. In perhaps the most important experiments, all three hits were tested for antiviral activity in SARS-CoV-2 assays. Piroxantrone and MMV1580853 did not display any detectable antiviral activity. Given their IC_50s_ against nsp15, Piroxantrone was too intrinsically cytotoxic to have detected an antiviral effect and MMV1580853 was probably too insoluble to be able to observe an antiviral effect. On the other hand, Exebryl-1 was shown to have SARS-CoV-2 antiviral activity in three different assays. One caveat for these experiments with Exebryl-1 was that the antiviral activity was sometimes close to the cytotoxicity (CC_50_), e.g. in Caco-2 cells. But clear separation of antiviral activity (EC50) from CC_50_ was observed in assays with Vero and Calu3 cells.

In developing our HTS, we found that Mn^2+^ but not Mg^2+^ was essential for endoU activity. Conversely, Mg^2+^ but not Mn^2+^ led to a significant (3-4°C) positive shift in the DSF T_m_ of apo nsp15. This positive T_m_ in DSF often implies that a ligand complexes and stabilizes a protein structure. However, in this case, Mg^2+^ (or Mn^2+^) is not unambiguously present in any of the deposited crystal structures including structures with substrates (PDB IDs: 6WLC, 6X4I, 7K0R) or transition state analogs (PDB ID: 7K1L). Thus, the reason for the DSF shift with Mg^2+^ is not understood. Manganese has long been known to be important for nsp15 endoU function [12, 14, 19]. But to date, the catalytic role of Mn^2+^ is unclear. One hypothesis is that Mn^2+^ may maintain the conformation of the RNA substrate during catalysis [15]. An alternative hypothesis, supported by a possible Mg^2+^ density in their nsp15 apo structure (PDB ID 6VWW), postulates that a metal binding site near the active site is necessary for active site stability during the cleavage reaction [13]. Further experiments, and ideally structures, are necessary to address the mechanistic role of divalent cations in the activity of nsp15.

CoV nsp15 forms a hexameric structure in solution and this hexameric structure has been hypothesized to be essential to the endoU function of nsp15 [12–14, 30, 31]. Indeed, the intermolecular bonds of the hexamer appear to stabilize the active site of nsp15 endoU. A recent preprint demonstrates nsp15 function in alpha coronavirus isolate, HCoV-229E, and chemical-genetic validation with a betulonic acid derivative (lead molecule **5h**) that inhibited replication of Coronavirus 229E at 0.6 μM EC50 [16]. It was shown that a nsp15 deficient coronavirus was much less sensitive to **5h** and selecting for mutants with **5h** led to mutations in the N terminus of nsp15 that map to residues important in the hexameric macromolecule of CoV nsp15. Unfortunately, the **5h** lead inhibitor had no activity against SARS-CoV-2 in Vero-E6 cells, and the inhibitor may be limited in utility to alpha coronaviruses and not to beta coronaviruses like SARS-CoV, SARS-CoV-2, and MERS-CoV.

The FRET-based nsp15 endoU screening of libraries for inhibitors identified many PAINS compounds that generated H_2_O_2_ in the presence of reducing reagents, such as DTT, as identified in the Amplex Red counter screen. All of these PAINS molecules that were positive for H_2_O_2_ generation in the Amplex Red assay led to a lower T_m_ in the DSF assay, suggesting the protein was destabilized in the presence of these molecules. Unfortunately, reducing reagents were required in the FRET assay, so we could not test the effect of PAINS molecules without H_2_O_2_ generation. We hypothesize that the generation of H_2_O_2_, and possibly other ROS, by these PAINS molecules inhibited enzyme activity by oxidative damage to the protein or substrate.

Our hits that could be reconfirmed in the non-fluorescent endoU assay included Exebryl-1 and Piroxantrone. While MMV1580853 exhibited inhibition of endoU cleavage in this assay, we were unable to test a sufficient range of concentrations to establish IC_50_ values, likely due to the compound’s solubility limitations in DMSO and assay buffer. MMV1580853, also known as CHEMBL3410452 (https://www.ebi.ac.uk/chembl/compound_report_card/CHEMBL3410452/), is an antibacterial lead compound developed for *anti-Staphyloccocus* activity (compound 17 in [32]). MMV1580853 inhibits bacterial isoprenoid biosynthesis, targeting undecaprenyl diphosphate synthase (UPPS) [32]. The relative insolubility and lack of potency of this compound limits its ability to be further refined, though it could serve as lead to improve potency and solubility. Piroxantrone is an anthrapyrazole anti-cancer investigational new drug that intercalates in DNA and has been used in phase 2 clinical trials for lung and breast cancer [33, 34]. However, Piroxantrone is highly toxic and unlikely to be a good lead for further development as a nsp15 inhibitor.

Our hit that was characterized to best inhibit nsp15 endoU activity, bind to nsp15, and express antiviral activity towards SARS-CoV-2 was Exebryl-1. Exebryl-1 was an analog synthesized to mimic natural products from the rain forest woody vine, *Uncaria tomentosa*, that had been shown to have β-amyloid inhibitory activity [35]. Exebryl-1 was developed by ProteoTech Inc. as a β-amyloid anti-aggregation molecule for Alzheimer’s therapy [35, 36]. It had been shown to disperse β-amyloid fibrils in vitro and to a decrease of neurofibrillary tangles in a mouse model of Alzheimer’s disease [36]. In previous studies, mice have been treated with 100 mg/kg per day of Exebryl-1 for 6 months [37], without evident toxicity, though we could not find records of formal pre-clinical safety testing. Exebryl-1 never made it to clinical trials, so the concentrations of what could be achieved in humans is unknown. However, the plasma levels likely would not reach therapeutic levels. A 100 mg/kg oral dose of Exebryl-1 in Sprague-Dawley rats only achieved 9 μM plasma concentrations at 1 h, with plasma levels below 4 μM by 4 h after dosing. By contrast, the antiviral effect in our assays required more than 10 μM concentrations. However, given its activity profile, while Exebryl-1 doesn’t seem to have immediate repurposing potential, it remains a prime lead for further medicinal chemistry optimization. The medicinal chemistry can be rationally driven using the predictions from our docking studies (**Fig 6**). While the endoU catalytic groove is the most apparent target binding site, our results also strongly suggest another druggable site deep inside the pore of the nsp15 hexamer. Ligand binding in the pore may have allosteric effect on the activity, although the mechanism of action is still unclear. The observation here highlights the implications of such unknown dynamics or long-range effects for protein function, which may not be fully captured by solved structures. Integration of multiple and complementary biophysical techniques can thus be beneficial.

The studies reported here suggest that nsp15 can be specifically targeted with inhibitors and nsp15 endoU inhibitors will have expected antiviral activity. Furthermore, as stated in the introduction, antiviral activity can be effective across the related coronaviruses, such that any inhibitors identified using the screening methodologies described here could be potential useful antivirals for other pathogenic coronaviruses which may emerge, in the future. To conclude, the results presented here supports expanded screening of SARS-CoV-2 nsp15 endoU against larger small-molecule libraries to identify potential inhibitors. Further studies are necessary to demonstrate whether simultaneous engagement of the innate immune system can amplify the antiviral effects of nsp15 inhibitors.

## Materials and Methods

### Libraries and chemicals

The ReFRAME drug repurposing library was provided by Calibr at Scripps Research (La Jolla, CA). The Pandemic Response Box and COVID Box libraries were provided by the Medicines for Malaria Venture (Geneva, Switzerland). The 5’6-FAM/dArUdAdA/3’-TAMRA oligonucleotide was purchased from IDT DNA Inc (Coralville, IA). The compounds DA-3003-1, Ceftazidime, μ-lapachone, and BVT-948 were purchased from Cayman Chemical (Ann Arbor, MI). The compounds Exebryl-1 and BN-82685 were synthesized by Los Alamos National Laboratory (Ryszard Michalczyk). The route of chemical synthesis for Exebryl-1 is outlined in **Fig S4**.

### Recombinant protein expression and purification

The SARS-CoV2 nsp15 gene sequence was codon optimized for *E. coli* expression and commercially synthesized and cloned into the pET-28a(+) vector with an N-terminal hexahistidine tag (GeneScript, Piscataway, NJ). The construct was transformed into Rosetta BL21(DE3) competent cells and expressed using autoinduction methods [38, 39] in 2L bottles in a LEX bioreactor at 25°C for 24 h followed by a drop in temperature to 15°C for 72 h. Cell pellets were harvested and lysed via sonication in buffer containing 25 mM HEPES pH 7, 500 mM NaCl, 5% glycerol, 30 mM imidazole, 0.025% sodium azide, 0.5% CHAPS, 10 mM MgCl_2_, 1 mM TCEP, 250 μg/mL AEBSF, and 0.05 μg/mL lysozyme. Lysate was incubated with 25 U/mL benzonase, centrifuged, and soluble supernatant was purified using HisTrap FF 5 mL immobilized metal ion affinity chromatography columns (GE Healthcare, New Jersey, USA) and eluted with 250 mM imidazole. Samples were fractionated on a Superdex 200 size-exclusion chromatography column (GE Healthcare) and an AKTA Explorer 100 System (GE Healthcare). Fractions were visualized via SDS-PAGE and concentrated using Amicon Ultra centrifugal filters. Samples were flash frozen in liquid nitrogen and stored at −80°C.

### Endoribonuclease FRET assay

To measure cleavage activity of nsp15 in a high-throughput manner, a previously reported fluorescence resonance energy transfer (FRET) assay was adopted [18, 19]. Briefly, a 4 nucleotide chimeric substrate was commercially synthesized (IDT DNA Inc., Coralville, IA) with a 5’ carboxyfluorescein (FAM) fluorophore and 3’ 5-carboxytetramethylrhodamine (TAMRA) quencher (5’6-FAM/dArUdAdA/3’-TAMRA) such that fluorescence is quenched until enzymatic cleavage of uridylate liberates the fluorophore. Recombinant nsp15 was diluted in assay buffer (50 mM Tris-HCl pH 7.5, 50 mM KCl, 5 mM MnCl_2_, 1 mM DTT) to a 25 nM concentration and incubated at ambient temperature with 0.5 μM FRET substrate in a final assay volume of 10 μL for 60 minutes. For the primary screen, 10 nL of the 10 mM ReFRAME library was pre-spotted onto black low-volume 384-well plates (Corning 3820) for a final assay concentration of 10 μM. For secondary and tertiary dose response assays, inhibitors were three-fold serially diluted and tested in duplicate at 10 μM and 100 μM top concentrations, respectively. Fluorescence was monitored on an Envision microplate reader (PerkinElmer, Waltham, MA) at excitation/emission wavelengths of 492 nm/518 nm. For baseline controls, nsp15 was denatured at 95°C for 5 min and added in place of intact enzyme; and for neutral solvent controls, DMSO was added in place of inhibitor. The dataset was normalized to the dynamic range of the assay and reported as percent inhibition. GraphPad Prism (GraphPad Software Inc) was used to plot dose response curves and calculate IC_50_ values.

### Differential scanning fluorimetry assay

Recombinant SARS-CoV2 nsp15 was diluted to 8 μM in assay buffer (50 mM Tris-HCl pH 7.5, 50 mM KCl, 5 mM MnCl_2_, and 1 mM DTT) containing 100 μM ligand and incubated at ambient temperature for 15 min. SYPRO^®^ Orange dye (Invitrogen) was diluted to a 5x concentration in assay buffer and 10 μL added to 10 μL of the enzyme/drug mixture in a 96-well plate. The plate was sealed and pulse centrifuged to consolidate and remove air bubbles. Fluorescence was continuously monitored at an excitation/emission wavelength of 300/470-570 nm using a StepOne Plus RT-PCR thermal cycler (Applied Biosystems, Foster City, CA) as the samples were heated from 25°C to 95°C at a ramp rate of 1°C/min. GraphPad^®^ Prism Software (GraphPad Software, San Diego, CA) was used to plot the first derivative of the rate of change of fluorescence (- d(RFU)/dT) versus temperature (°C), and the local peak minima was reported as the protein melting temperatures (T_m_).

### Amplex Red assay

To assess the redox cycling potential of compounds in the presence of the reducing agent, dithiothreitol (DTT), a fluorometric assay (Invitrogen, Carlsbad, CA) was employed that uses the 10-acetyl-3,7- dihydroxyphenoxazine (Amplex Red) reagent to detect the generation of hydrogen peroxide. In black 384 plates, inhibitors were diluted to a 100 μM concentration using kit provided assay buffer along with varying concentrations of DTT (0, 0.2, and 2 mM). Fluorescence was measured on an Envision microplate reader (Perkin Elmer, Waltham, MA) at excitation/emission wavelengths of 531/595 nm to obtain background fluorescence levels. An equal volume containing 100 μM Amplex Red and 0.2 U/mL horseradish peroxidase was then added to the plates and allowed to incubate at ambient temperature for 15 min, protected from light. Plates were read a final time and corrected fluorescence values were compared to DMSO control. DA-3003-1, a quinone-based compound with high redox cycling potential, was included as a positive control.

### Native MS

Recombinant nsp15 was buffer exchanged into 100 mM ammonium acetate (pH adjusted to 7.5 using ammonium hydroxide) using size exclusion spin columns (Zeba Micro Spin Desalting Columns, 7k MWCO, 75 μL) following manufacturer’s protocol. The protein solution was then diluted to 5 μM, supplemented with 10 βM Mn(II) acetate, and mixed with ligands at desired concentrations. Since the tested ligands were dissolved in DMSO, supplemental DMSO was added to the control and all concentrations of Exebryl-1 below 10 μM such that the minimum amount of DMSO was 0.1% by volume (same as the 10 μM Exebryl-1 sample). At low concentrations, DMSO is known to shift the charge state distribution (higher or lower) of proteins in nanoelectrospray [40]. In this case, a charge-reducing effect was observed for nsp15 with DMSO, and the addition of supplemental DMSO corrected this effect for the control and low concentrations of Exebryl-1 (<10 μM Exebryl-1). Native MS was performed using a Waters Synapt G2s-i ion mobility time-of-flight (TOF) mass spectrometer (Waters, Milford, MA). A minimally invasive SID device (kindly provided by Prof. Vicki Wysocki’s group [41]) was installed via custom modification of the DRE (dynamic range enhancement) lenses (after the mass isolation quadrupole). Samples were introduced in positive mode with the static nanoelectrospray source via in house pulled borosilicate glass capillaries (Sutter Instrument, model P-1000, capillary item # BF100-78- 10). Electrospray voltage (0.6-1 kV) was applied via a Pt wire inserted to the capillary and in contact with the protein solution. Source temperature was set to 30°C without any gas flow, cone was at 150 V. Trap gas flow was set to 4-8 mL/min to maximize signal during SID/CID. Data were collected in TOF, sensitivity mode. Sodium iodide clusters were used for mass calibration. A quad profile set to 6000, 8000, and 10000 was used to remove monomers prior to gas-phase activation, while maintaining high signal levels of the nsp15 hexamer. Collision voltage in CID or SID was applied in the trap traveling wave ion guide. To perform SID in the modified instrument, the TOF collector was set 10 V lower than trap collision voltage. The TOF stopper was set to ½ of the collector voltage. Signal were typically averaged over 5-10 min for each spectrum.

Spectra were analyzed manually in MassLynx v4.1 (Waters, Milford, MA). Mass deconvolution for monomers and hexamers in the CID/SID spectra was performed in UniDec v4.3.0 [42]. The mass profiles (**Fig 6D-E**) are heterogenous from buffer adducts and PTMs, making it difficult to directly read out the binding stoichiometry due to overlapping profiles. We used the “double deconvolution” function in UniDec [43] to fit the ligand bound mass profiles using apo mass profiles as “kernel”. This analysis essentially combines the species from buffer adducts/PTMs, and yielded the relative intensities of the released monomers with different numbers of bound Exebryl-1. Peak intensities were then used to calculate the percent relative abundance in **Fig 5F** and average ligand numbers per 1mer in **Fig S1**.

### Molecular docking simulations

Molecular docking simulations were carried out with AutoDock Qvina[28, 29] by restricting search space to the receptor binding pocket of the nsp15 crystal structure (PDB ID: 6XDH). We used our automated workflow for docking which converts compound SMILES into PDBQT/PDB format using RDKit and OpenBabel [44]. All other parameters of the Qvina code were kept as default and every bond in the compound was allowed to rotate freely with the protein receptor rigid. Moreover, Qvina uses multiple CPU cores [45] which reduces its running time and it was set up on a high-performance computing cluster which generates a file for each compound with molecular coordinates for up to 20 poses along with the Gibbs free energy and the scoring function for each binding model [46]. Each compound was docked in the specified binding pocket of nsp15 monomer and was sorted by maximum binding score for analyzing potential binding sites.

### Mammalian cytotoxicity assay

Lead compounds were tested in vitro against two mammalian cell lines to evaluate cytotoxicity. CRL- 8155 human lymphocyte and HepG2 human hepatocyte cells (ATCC, Manassas, VA) were seeded in 96-well plates and incubated at 37°C for 48 h in the presence of test compound (serial-2 dilutions, in triplicate). At the end of the incubation period, cells were visually assessed before alamarBlue^™^ (Thermo Fisher, Waltham, MA), a resazurin-based cell viability reagent which measures metabolic activity, was added to the plates and fluorescence measured on a BioTek FLx-800 microplate reader (BioTek Instruments, Winooski, VT). Fluorescence signals resulting from cell viability changes were compared with control wells to calculate 50% cytotoxic concentrations (CC_50_) values.

### In vitro RNA cleavage assay

The RNA cleavage activity of nsp15 was assessed using a 31 nucleotide (nt) RNA oligonucleotide substrate (IDT DNA) consisting of a string of rA bases punctuated by a single rU (rArArArArArArArArArArArArArArArArArArArArUrArArArArArArArArArA) such that cleavage results in 20 nt and 10 nt fragments. Test compounds were serially diluted in DMSO and pre-incubated with 0.78 μM nsp15 in assay buffer consisting of 50 mM Tris-HCl pH 7.5, 50 mM KCl, 5 mM MnCl_2_, and 1 mM DTT for 10 min. The reaction was initiated with the addition of 500 ng RNA substrate and allowed to proceed for 2 h at room temperature, with a final DMSO concentration of 1%. The reaction products were electrophoresed on a 15% polyacrylamide gel and run at 100 V for 2 h at 4°C. Gels were stained with SYBR green II (Thermo Fisher) for 30 min and visualized on a Bio-Rad Gel Doc (Bio-Rad Laboratories, Hercules, CA). Densitometry was performed using Bio-Rad Image Lab software. The ratio of cleaved vs uncleaved substrate was calculated and normalized to nsp15 (0%) and DMSO controls (100%).

### In vitro CPE assay in Vero cells

The antiviral activities of test compounds were evaluated in Vero 76 cells. Eight dilutions were tested and the effective antiviral concentration determined by regression analysis. The toxicity of the test compounds were determined in parallel. CPE was determined by microscopic observation of cell monolayers as well as uptake of neutral red dye.

### In vitro VYR assay in Caco-2 cells

As a follow-up to the CPE assay, a viral yield reduction assay was performed to evaluate the ability of test compounds to inhibit virus production in Caco-2 cells. Virus was introduced to cells containing varying dilutions of test compound. After an incubation period, viral titer was determined by endpoint dilution in 96-well microplates. Eight dilutions of test compound were assayed, and effective antiviral concentrations were determined by regression analysis.

### In vitro antiviral assay in Calu3 cells

Ten thousand Calu-3 cells (HTB-55, ATCC) grown in Minimal Eagles Medium supplemented with 0.1% non-essential amino acids, 0.1% penicillin/streptomycin, and 10% FBS were plated in 384 well plates. The next day, 50 nL of drug suspended in DMSO was added to assay plates as an 8-point dose response with three-fold dilutions between test concentrations in triplicate, starting at 200 μM final concentration. The negative control (0.2% DMSO, n=32) and positive control (10 μM Remdesivir, n=32) were included on each assay plate. Calu3 cells were pretreated with controls and test drugs (in triplicate) for 2 h prior to infection. In BSL-3 containment, SARS-CoV-2 (isolate USA WA1/2020) diluted in serum free growth medium was added to plates to achieve an MOI=0.5. Cells were incubated continuously with drugs and SARS-CoV2 for 48 h. Cells were fixed and then immunostained with anti-dsRNA (mAb J2) and nuclei were counterstained with Hoechst 33342 for automated microscopy. Automated image analysis quantified the number of cells per well (toxicity) and the percentage of infected cells (dsRNA+ cells/cell number) per well. SARS-CoV-2 infection at each drug concentration was normalized to aggregated DMSO plate control wells and expressed as percentage-of-control (POC=% Infection _sample_/Avg % Infection _DMSO cont_). A non-linear regression curve fit analysis (GraphPad Prism 8) of POC Infection and cell viability versus the log10 transformed concentration values was used to calculate IC_50_ values for Infection and CC_50_ values for cell viability. Selectivity index (SI) was calculated as a ratio of drug’s CC_50_ and IC_50_ values (SI = CC_50_/IC_50_).

## Acknowledgements

The authors would like to thank: all of the members of the Seattle Structural Genomics for Infectious Diseases (SSGCID.org) many of whom contributed to science and infrastructure supporting this project; Dr. Arnab Chatterjee, Emily Chen, Mitchell V. Hull, and the Compound Management Group at Calibr, a division of The Scripps Research Institute, for supporting the organization of the library distribution and compound follow-up.; the Medicines for Malaria Venture Open program (https://www.mmv.org/mmv-open) for access to their compound libraries and for sending follow-up compounds, and particularly the assistance and advice of Drs. Kirandeep Samby and Tim Wells; Prof. Vick Wysocki, Dr. Dalton Synder, and Benjamin Jones (Ohio State University) for assistance with SID and Prof. Michael Marty (University of Arizona) for assistance with UniDec; and, Dr. David C. Schultz and the University of Pennsylvania High-throughput Screening Core for supporting the in vitro anti-SARS-CoV-2 studies in Calu-3 cells. Part of the research was conducted at the Environmental Molecular Sciences Laboratory (grid.436923.9), a national scientific user facility sponsored by the U.S. Department of Energy’s Office of Biological and Environmental Research (BER) program located at Pacific Northwest National Laboratory (PNNL). Battelle operates PNNL for the U.S. Department of Energy under contract DE-AC05-76RLO-1830. The University of Washington has utilized the non-clinical and pre-clinical services program offered by the National Institute of Allergy and Infectious Diseases. Following the open access policy for screening the Calibr/Scripps ReFRAME library, these data have been deposited into a freely accessible database at reframedb.org.

## Supporting information

**Fig S1.**
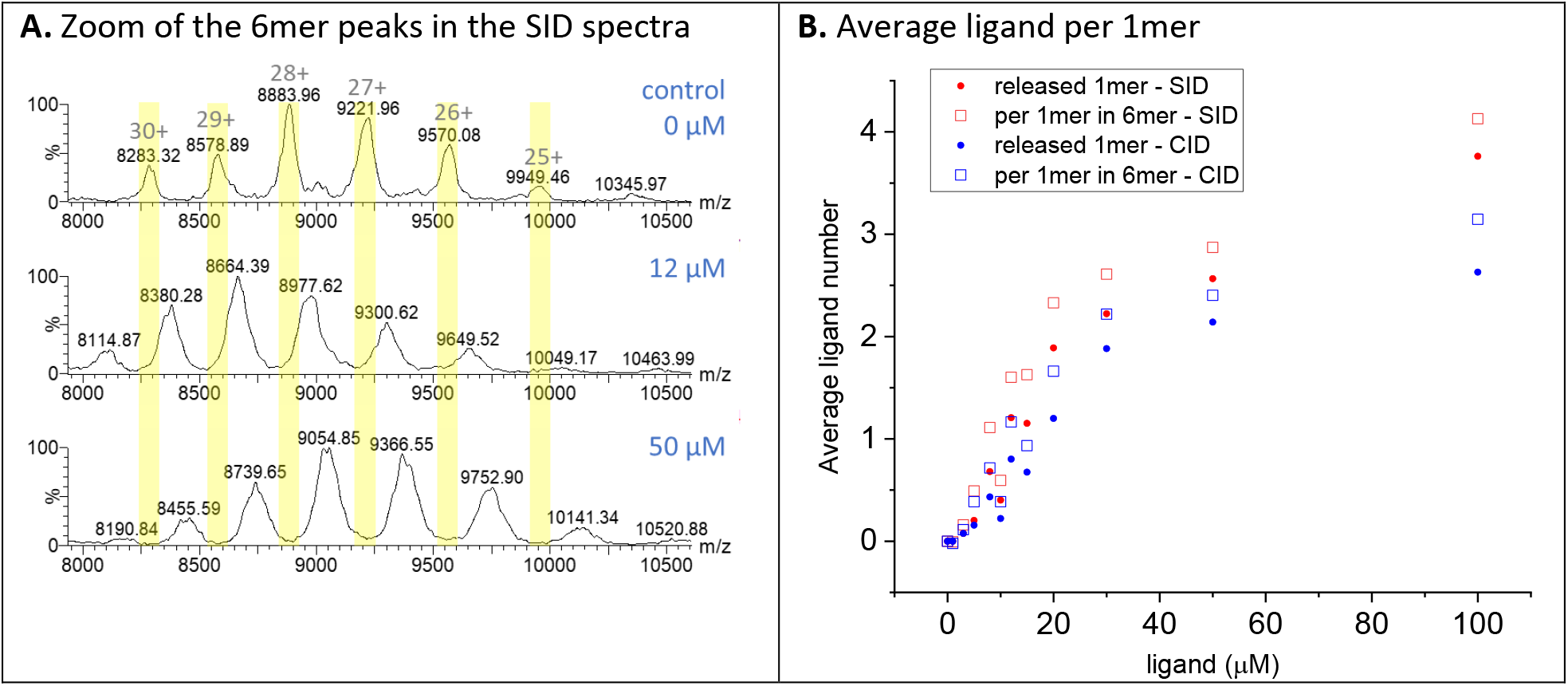
Exebryl-1 mass addition on the nsp15 hexamer. **A)** Zoom-in view of the 6mers peaks from SID spectra at different Exebryl-1 concentration. Significant mass increase was detected as ligand concentration increased, as seen by the shift of all peaks to the right. **B)** Average ligand per 1mer based on the data from released 1mers (filled circles) and mass shift on the 6mers (open squares) from SID (red) and CID (blue). The total mass shifts on the 6mer was divided by the mass of Exebryl-1 to yield the total number of ligands. Then the number was further divided by 6 to yield the average ligand per 1mer, assuming all ligands are symmetrically distributed within the 6mer. The mass shifts correlate well with the mass shift on 1mers, suggesting Exebryl-1 binds to all 1mers in the complex symmetrically. The systematic higher values calculated from 6mers than from 1mers are likely due to additional nonspecific salt/solvent addition to the 6mers and potential loss of weakly associated exebryl-1. CID consistently showed lower ligand numbers on both 1mers and 6mers, especially at increased ligand concentration.

**Fig S2.**
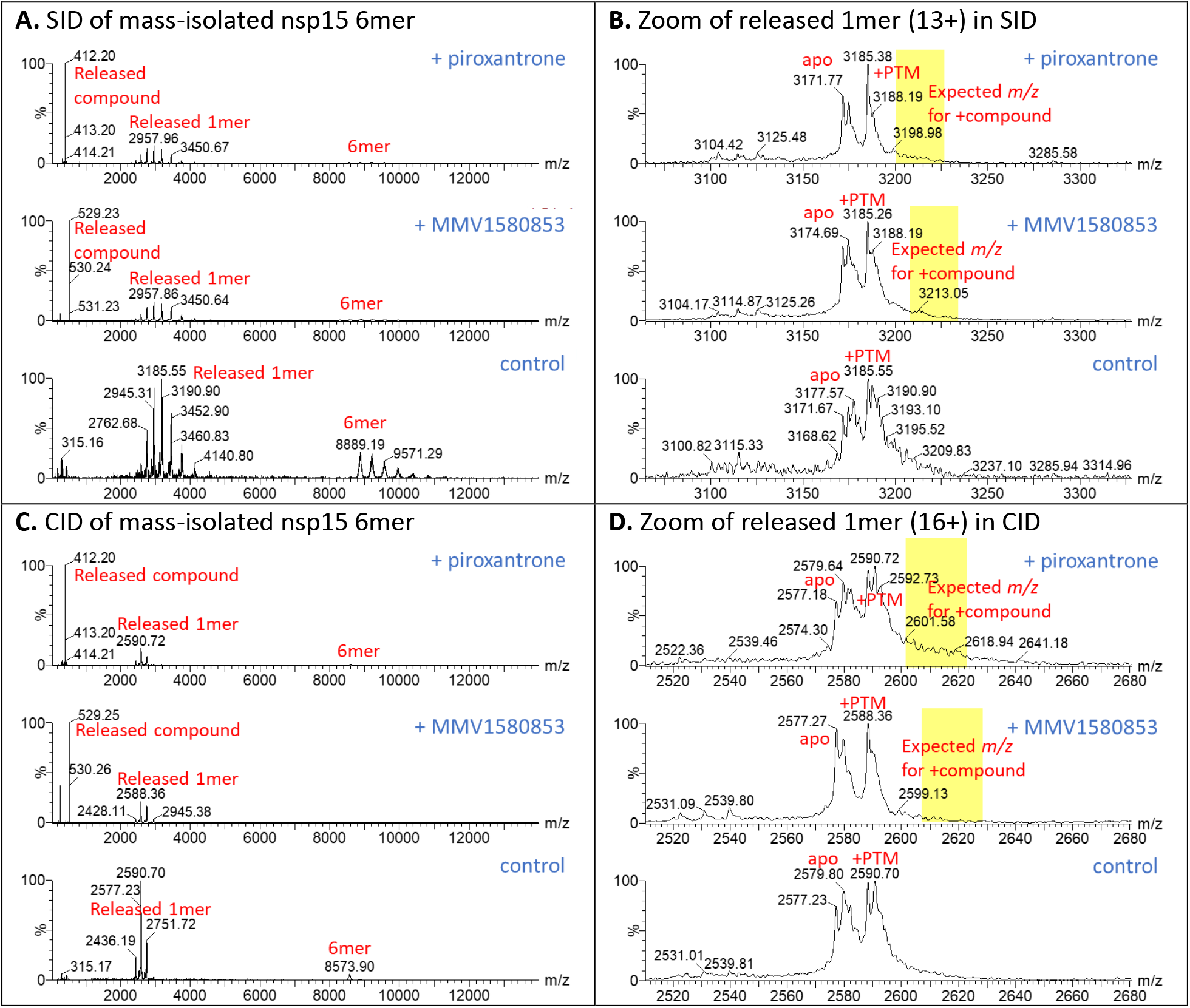
Native MS analysis of nsp15 binding to Piroxantrone and MMV1580853. **A)** Full views of 90 V SID for mass isolated nsp15 6mer. **B)** Zoom view of the released 1mer in the SID spectra. **C)** Full views of 100 V CID for mass isolated nsp15 6mer. **D)** Zoom view of the released 1mer in the CID spectra. The top and middle panels in each panel are the spectrum with compounds added (Piroxantrone and MMV1580853, respectively), the bottom panel is control. The same experimental methods used for exebryl-1 binding was applied here. Nsp15 at 5 μM concentration in 100 mM ammonium acetate (pH 7.5) was mixed with 10 μM Mn(II) acetate, and 10 μM compound. The compound was diluted from 10 mM stock in DMSO. For control, pure DMSO was used as stock. No significant amount of piroxantrone or MMV1580853 can be detected in the released 1mers in both SID and CID at 10 μM compound concentration. However, the compounds were released from the 6mer upon activation, as shown by the strong signal at low *m/z* (412.2 and 529.2 for protonated piroxantrone and MMV1580853, respectively). The data suggest these compounds weakly bind to nsp15.

**Fig S3.**
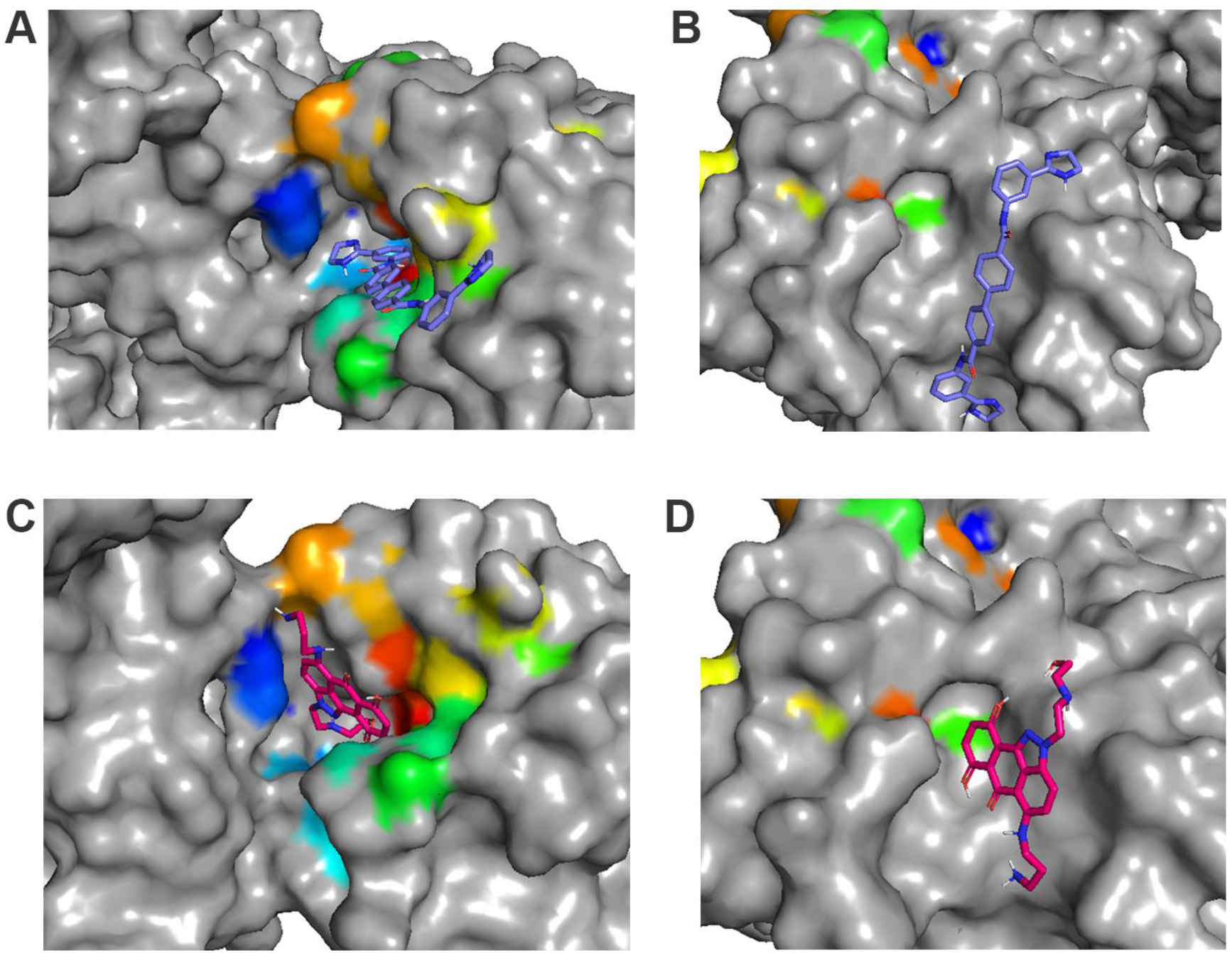
Molecular docking of MMV1580853 and Piroxantrone. A) MMV1580853 binding mode 1 in a deep pocket formed between the C-terminal endoU and N-terminal oligomerization domains and oriented within the pore structure of the catalytically active hexamer. B) MMV1580853 binding mode 2 in the endoU catalytic site (relatively less stable by 1.1 kcal/mol, lower binding affinity as compared to mode 1). C) Piroxantrone binding mode 1 within the pore structure. D) Piroxantrone binding mode 2 in the endoU catalytic site (relatively less stable by ~0.6 kcal/mol, lower binding affinity as compared to mode 1). MMV1580853 and Piroxantrone appear to bind on the surface in both binding modes rather than buried within the binding pockets.

**Fig S4.**
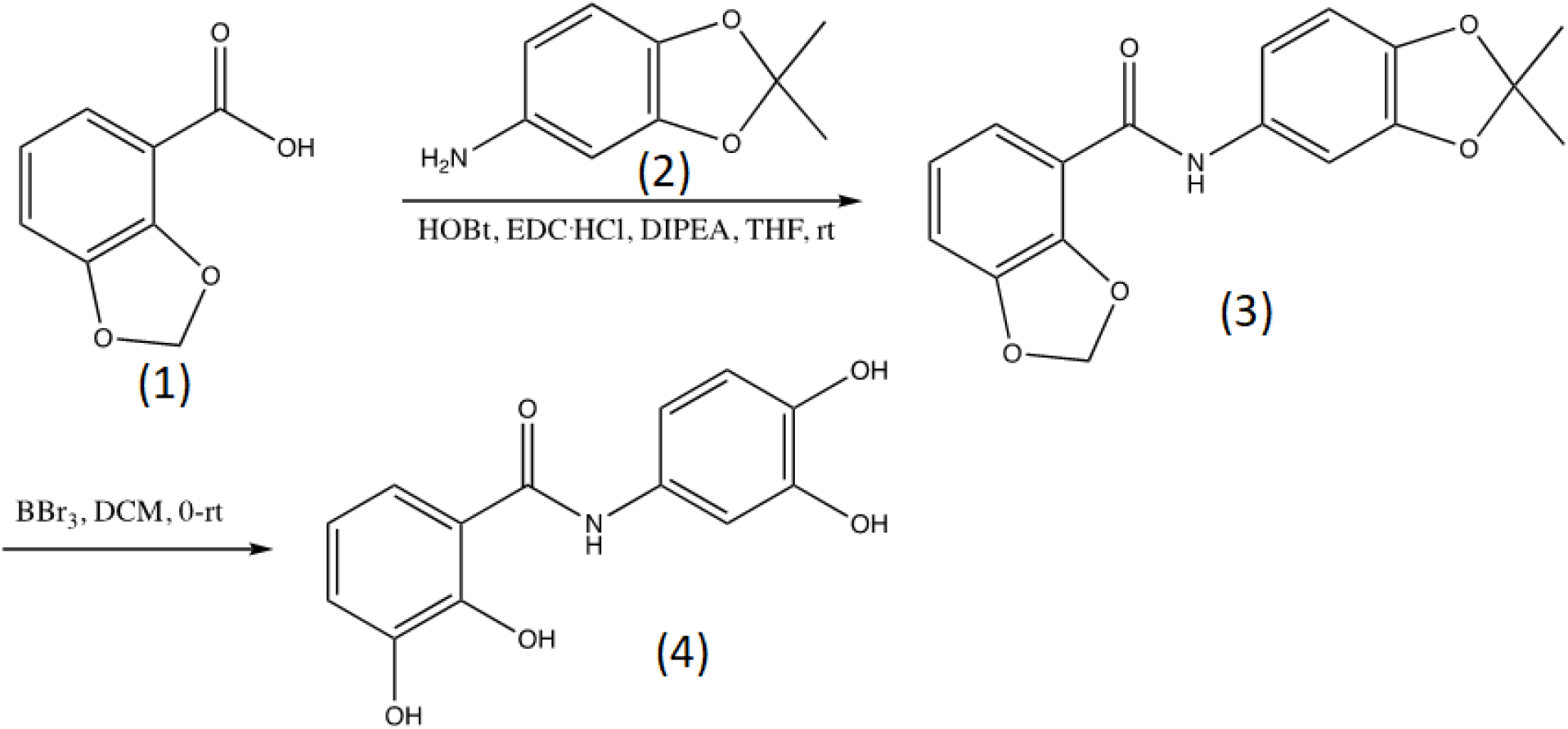
Chemical synthesis of Exebryl-1. A brief description of the synthesis is as follows: to a suspension of protected benzoic acid **(1)** (0.64 g, 3.85 mmol) in THF (10.0 mL) was added hydorxybenzotriazole (HOBt, 0.52 g, 3.85 mmol), 1-ethyl-3-(3-dimethylaminopropyl)carbodiimide (EDC.HCl, 0.738 g, 3.85 mmol) and diisopropylethylamine (DIPEA, 1.83 mL, 10.5 mmol) at room temperature and stirred for 30 min. To this now clear solution the protected aniline **(2)** was added (0.48 g, 2.9 mmol) and the mixture was stirred at room temperature overnight. The solvent and excess reagent were removed under reduced pressure and the residue was purified on silica gel with 10% EtOAc/hexanes to provide 0.56 g (62%) of the intermediate **(3)** as pinkish solids. To a solution of the intermediate product **(3)** (0.56 g, 1.69 mmol) in DCM (17 mL) was added BBr3 (0.65 mL, 6.76 mmol) at 0°C, the mixture was allowed to warm to room temperature and stirred overnight. Methanol was added slowly until the suspension became clear. The solvents and excess reagent were removed under reduced pressure and the residue was purified on silica gel with 60% EtOAc/hexanes followed by 10% MeOH/DCM to provide the product with pink color. The material was suspended and stirred in DCM, filtered and washed with DCM to provide the final product as white solids (0.3 g, 68%).

